# FrameRate: learning the coding potential of unassembled metagenomic reads

**DOI:** 10.1101/2022.09.16.508314

**Authors:** Wang Liu-Wei, Wayne Aubrey, Amanda Clare, Robert Hoehndorf, Christopher J. Creevey, Nicholas J. Dimonaco

## Abstract

**Motivation:** Metagenomic assembly is a slow and computationally intensive process and despite needing iterative rounds for improvement and completeness the resulting assembly often fails to incorporate many of the input sequencing reads. This is further complicated when there is reduced read-depth and/or artefacts which result in chimeric assemblies both of which are especially prominent in the assembly of metagenomic datasets. Many of these limitations could potentially be overcome by exploiting the information content stored in the reads directly and thus eliminating the need for assembly in a number of situations.

**Results:** We explored the prediction of coding potential of DNA reads by training a machine learning model on existing protein sequences. Named ‘FrameRate’, this model can predict the coding frame(s) from unassembled DNA sequencing reads directly, thus greatly reducing the computational resources required for genome assembly and similarity-based inference to pre-computed databases. Using the eggNOG-mapper function annotation tool, the predicted coding frames from FrameRate were functionally verified by comparing to the results from full-length protein sequences reconstructed with an established metagenome assembly and gene prediction pipeline from the same metagenomic sample. FrameRate captured equivalent functional profiles from the coding frames while reducing the required storage and time resources significantly. FrameRate was also able to annotate reads that were not represented in the assembly, capturing this ‘missing’ information. As an ultra-fast read-level assembly-free coding profiler, FrameRate enables rapid characterisation of almost every sequencing read directly, whether it can be assembled or not, and thus circumvent many of the problems caused by contemporary assembly workflows.

**Availability:** https://github.com/NickJD/FrameRate

## 1 Introduction

The current availability and throughput of DNA sequencing technologies have drastically reduced the time and cost required for the sequencing of large, complex, and niche genomes and metagenomes from increasingly diverse environments [Land et al., 2015, Goodwin et al., 2016]. Contemporary research and computational advances in *de novo* genome assembly and annotation have struggled to keep pace with this revolution. Among their many limitations, genome assembly tools fail at assembling metagenomic samples, and often produce chimeric contigs that are not representative of what exists in nature [Alneberg et al., 2018, Chen et al., 2020]. Although techniques for the assembling and annotation of metagenomes are improving, the speed at which metagenomic datasets can be assembled and studied is still too slow when compared to the rate that new metagenomic data are being produced [Lapidus and Korobeynikov, 2021]. In addition to this, even on single genomes, genome annotation continues to be a developing field, with many known and unknown shortcomings that have yet to be addressed [Dimonaco et al., 2021].

There exists a class of assembly-free tools that aim to gain biological insight from (meta)genomic DNA sequencing reads. These tools are most often k-mer or homology based and require large pre-computed databases, and when applied to large sequencing projects, are often only tractable on large computational clusters [Bağci et al., 2021]. Such tools are often available with ‘reduced’ databases for lightweight analysis, there is nonetheless a reduction in utility as a result of this compromise [Wood et al., 2019]. There has recently been a trend to undertake read-level analysis on ‘local machines’, without the need for large computing infrastructure. For example, Kraken, which uses species-specific k-mer frequencies to assign taxonomy to both unassembled and assembled reads, has recently been reworked to no longer require large amounts of local storage and system memory to perform a ‘complete’ classification [Pockrandt et al., 2022]. Another such tool, Plass [Steinegger et al., 2019], specifically aims to partly circumvent genome assembly and gain protein-level insight directly from DNA sequencing reads. Plass performs protein-level assembly by assembling Open Reading Frames (ORFs) identified directly from unassembled DNA reads. However, since the first stage of Plass is to identify ORFs, which is known to be a difficult and error-prone [Dimonaco et al., 2021], it must filter out incorrect coding frames based on amino acid and dipeptide frequencies after the contigs are assembled.

While in theory for a sequence of *n* amino acids there are 20^*n*^ possible amino acid sequences, which implies a vast diversity, in practice there exists a number of biological and chemical constraints in both the DNA and amino acid sequence of the protein coding regions of a genome that significantly reduce this space [Alberts et al., 2002, Staden, 1984, Gribskov et al., 1984, Fickett, 1982]. The identification and exploitation of these patterns or rules have led to a number of advances in computational techniques such as the AlphaFold platform, which uses rules learned from the Protein Data Bank of protein structures [wwPDB consortium, 2019] to predict structure [Jumper et al., 2021]. Additionally, as the vast majority of a prokaryotic genome is protein coding, one could expect that the majority of DNA sequencing reads from prokaryote-rich environments would belong to a protein coding gene. Therefore, if these reads could be directly translated into their amino acid counterpart, without the requirement for assembly or ORF prediction, many hurdles inherent to genomic annotation could be overcome. However, this is not a straightforward task, since each read can potentially be translated into 6 different amino acid sequences (3 frames on each strand), among which only one (or a few, in the case of overlapping or closely positioned genes) is the true amino acid counterpart of the read.

Therefore, we hypothesise that the intrinsic biochemical rules that govern the protein-coding space can be learnt by an algorithm to identify true coding frames. To do this, we developed FrameRate, a convolutional neural network (CNN) that can quickly identify the correct coding frame(s) of DNA sequencing reads without assembly. Trained on a curated dataset of existing CDS sequences of thousands of bacterial genomes in over 100 genera, FrameRate was first tested for its ability to predict the correct coding frames in diverse gene families and further validated through alignments to existing protein databases such as Swiss-Prot and the Hungate Collection. Lastly, we applied FrameRate on a metagenomic sample and used the predicted coding frames directly for functional profiling and compared them against the functional profiles obtained from the CDS genes predicted from the metagenome assembly. We show that FrameRate, as a versatile ultra-fast read-level coding frame predictor, allows for rapid functional characterization of an entire metagenomic sample, even for reads that cannot be assembled.

## 2 Methods

### 2.1 Training Data

The 46^th^ release of Ensembl Bacteria [Yates et al., 2022] was selected as the training data for FrameRate. Highly fragmented genomes were removed by filtering out those with more than 5 contigs (not counting plasmids). Then, genera that had less than 5 genomes were removed. This resulted in 6,223 genomes which were from 1,417 species and 179 genera (see Supplementary Table 1). For each genome, the complete set of CDS DNA sequences were downloaded for a combined total of 21,506,454 sequences. Each DNA sequence was translated into its respective amino acid sequence using the universal coding table. The universal coding table is used during this training stage because when we later apply FrameRate to unassembled reads, the correct coding table will not be known.

CD-HIT [Fu et al., 2012] was used to reduce the redundancy of the translated amino acid sequences by clustering them. All CD-HIT clustering was executed with these parameters; a sequence identity of 80% and shorter-sequence length difference of 80%, and the option to cluster the sequences to their most similar cluster (see Subsection ‘Gene Clustering with CD-Hit’ for the full command). CD-HIT chooses one representative sequence from each cluster, resulting in a reduced set of 3,931,852 sequences. 79,684 (0.37%) amino acid sequences were pre-filtered out by CD-HIT for containing a stop codon used to code for an amino acid.

This collection of amino acid sequences constitutes a large resource of sequences with and without experimental validation. Therefore, to validate these, we next used eggNOG-mapper [Cantalapiedra et al., 2021] to assign 3,449,532 of these amino acid sequences a gene family from the EggNOG database [Huerta-Cepas et al., 2019]. Each amino acid sequence with a hit to an eggNOG gene family was converted into 5 other potential frames, representing the true negatives in our data.

Then a multi-FASTA format file containing 20,697,192 sequences (3,449,532 coding and 17,247,660 non-coding frames) was built as the input to model training from the six frames for each CDS gene.

The training dataset was analysed using CD-Hit to evaluate the shared sequence identity between coding and non-coding frames. Out of the 20,697,192 coding and non-coding frames, only 53,950 formed into clusters with more than one sequence (forming 26,679 clusters). Most of these clusters were formed from non-coding sequences and very few were a mix of coding and non-coding. Of these 26,679 clusters: 15,683 clusters were formed from non-coding frames only, 9,972 clusters were formed from coding frames only, and just 1,024 clusters were formed from a mix of both. These clusters were also very small with a mean number of sequences for each being *∼*2 and a maximum of 9. It could have been expected that the number of non-coding only frames would be equal to that of 5 times the number of coding only frames. However, as the CD-HIT clustering was conducted on the amino acid sequences from the 6 frames from each CDS gene, the variability inherent to the codon to amino acid translation offers a vast potential for possible amino acid sequences of the 5 non-coding frames. Additionally, the small number of mixed clusters could themselves in-part be explained by error in the CDS annotations and the presence of overlapping genes.

### 2.2 Constructing the model

To learn to differentiate the coding and non-coding frames of a protein coding sequence, FrameRate employs a Convolutional Neural Network (CNN) model. The model takes as input an amino acid sequence between 50-75 residues in length. The amino acid sequence is represented as a one-hot embedding matrix of dimension (20, 75) where the first dimension correspond to the 20 common amino acids and the second dimension is the sequence length. Sequences shorter than 75 were padded with vectors of zeros at the end, representing absence of any amino acid.

The hyperparameters investigated in this study were: convolution filter size, number of filters, MaxPooling layer size, and the number of neurons in the fully connected layers. These parameters were optimized through extensive parameter search and then fixed throughout all of the results in this paper. The convolutional layers of the model consist of 32 convolutional filters of dimensions 2, 4, …, 16 respectively, totalling 256 filters. The filters were followed by a MaxPooling layer of dimension 1 and a fully connected layer of 32 units.

Instead of randomly splitting for training, testing, and validation datasets, the dataset was split at the gene family level, to prevent overrepresentation of larger gene families in our dataset. We grouped gene families based on their sizes, i.e. the number of genes belonging to them: less than 10 genes, 10 - 100, 100 - 1,000, 1,000 - 5,000, and more than 5,000 genes. The splitting of the dataset is then performed by sampling the same number of genes from each size group, for training, validation and testing sets respectively.

### 2.3 Frame prediction for reads

Given a (meta)genomic readset in the fasta file format, FrameRate classifies the reads into their correct frames in 2 stages: (1) converting the DNA sequencing reads into all 6 potential amino acid sequences and applying stop codon filtering, (2) apply the trained model on each amino acid sequence (frame) independently.

Conversion of each of its 6 possible amino acid sequence frames is done using the universal codon table. For each frame, the longest region uninterrupted by stop codons is selected and then truncated to a maximum of 75 amino acids from the 3’ end or stop codon. It has long been hypothesised that the over-representation of out-of-frame stop codons within CDS sequences is a defence against the negative effects of frameshift mutation and ribosomal slippage [Tse et al., 2010]. Therefore, the frames that have stop positions between 10 and 65 (first and last 10 amino acids) are removed. This allows for a minimum of 55 amino acids if a sequence has two stops at the ends of that range, which is still larger than the small-protein length most often used in the literature [Bartholomäus et al., 2021].

### 2.4 Metagenomic sequence data, assembly and protein prediction

Due to their diversity and thus difficulty for assembly, ruminant metagenomic datasets are often subject of metagenomic quality and benchmarking studies [Roumpeka et al., 2017]. Accordingly, the well-studied and species-rich metagenomic dataset of sheep ruminant methane yield[Shi et al., 2014] was selected to evaluate FrameRate. Specifically, the samples SRR873595_1/2 were selected from NCBI project no. PRJNA202380. Each FASTQ file consisted of 224,630,639 reads and Trimmomatic [Bolger et al., 2014] was used to remove adapters and trim the reads according to default quality control parameters. This resulted in the two FASTQ files with 214,611,607 reads each (see Supplementary Subsection Read Trimming for more detail). PandaSeq [Masella et al., 2012] was then used to pair-end join the reads from the two trimmed FASTQ files using default parameters. A single FASTA file was produced with 186,941,580 paired-end reads with the median length of 224 nt.

The paired-end reads were then used as the input data for the metagenomic assembly by MEGAHIT [Li et al., 2015] using default parameters. Of the 2,762,998 contigs made by MEGAHIT, 539,022 remained after removing those that were shorter than 1,000 bp. 1,647,050 CDS genes were identified in these contigs using Prodigal [Hyatt et al., 2010] with default parameters. Further details of the reads, contigs and CDS genes discussed in this section are in Supplementary Table 2.

### 2.5 Preparing Data for Comparisons

In this study, a number of different comparisons were conducted between the CDS genes predicted by Prodigal and the unassembled DNA reads from the rumen metagenomic sample. We formed three collections of reads from which to make these comparsions.

The first (“General”) consisted of a collection of 10 million reads which were randomly subsampled from the complete set of 186,941,580. This collection is representative of a typical use case for FrameRate, to classify reads from a metagenome, and will include reads that are not part of CDS genes.

Next we form two further collections to specifically explore the reads that align to the Prodigal CDS genes in the assembly (“CDS-Aligned reads”), and only the reads that did not align to any part of the assembly (“Unaligned reads”). Each of these collections was also randomly downsampled to a size of 10 million.

To create the CDS-Aligned and Unaligned read collections, the reads were aligned to the MEGAHIT metagenomic assembly with Bowtie2 [Langmead and Salzberg, 2012]. Using IntersectBed [Quinlan and Hall, 2010], the reads which aligned (or intersected) with the GFF file produced by Prodigal were reported as a BAM file and then converted to a FASTA file with samtools containing 132,254,283 of the original 186,941,580 (70.75%) pair-ended reads. Next, the same process was used to extract reads that did not align to the assembled contigs, resulting in 50,412,461 reads which were not assembled with MEGAHIT.

Subsampling to 10 million reads for each of these collections was done using Seqtk [HengLi, 2018]. These three ‘shallow’ subsamples were classified with FrameRate and functionally profiled, and the results compared to those from the metagenome CDS genes predicted by Prodigal. Figure 2 shows the production of the read collections and the application of FrameRate to these sets of reads and to the CDS genes found by Prodigal in the metagenome assembly.

**Figure 1:**
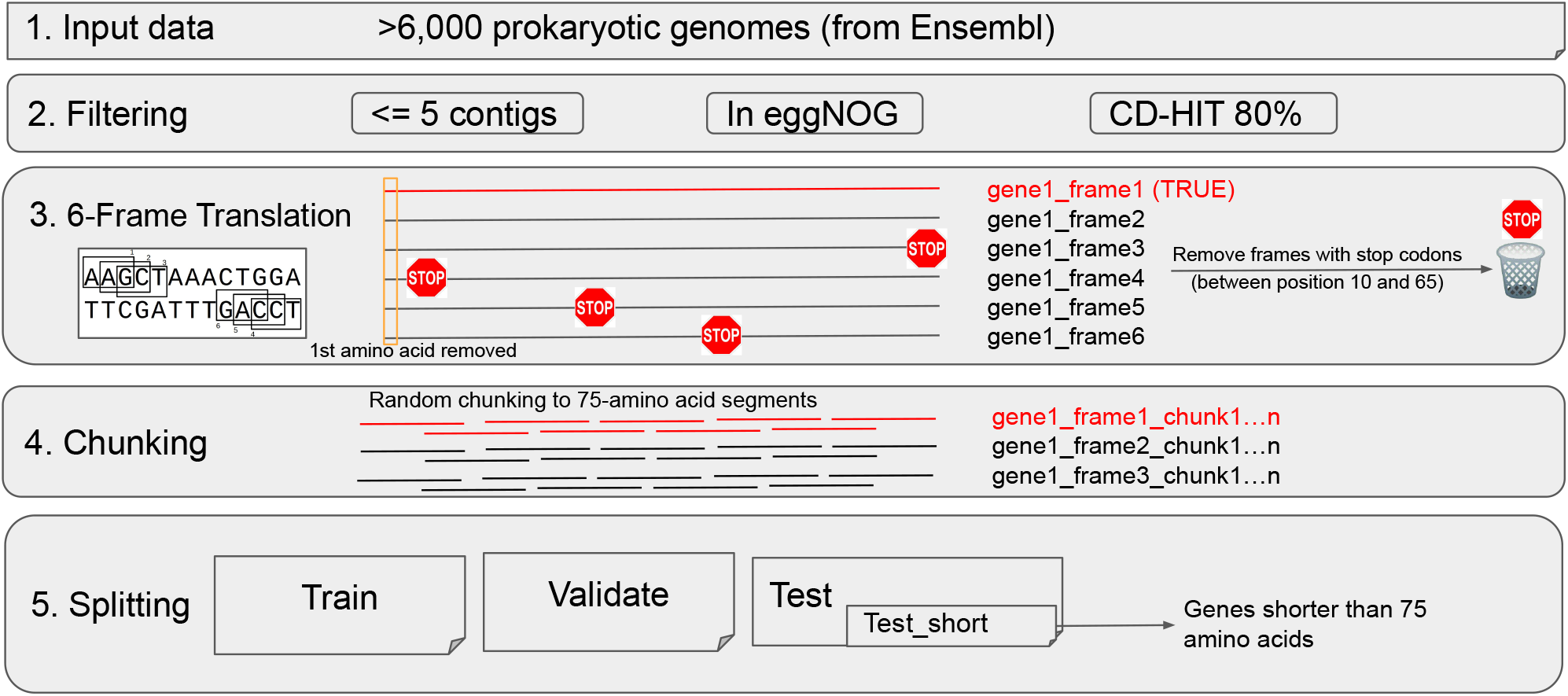
Overview of the data preprocessing for training FrameRate.

**Figure 2:**
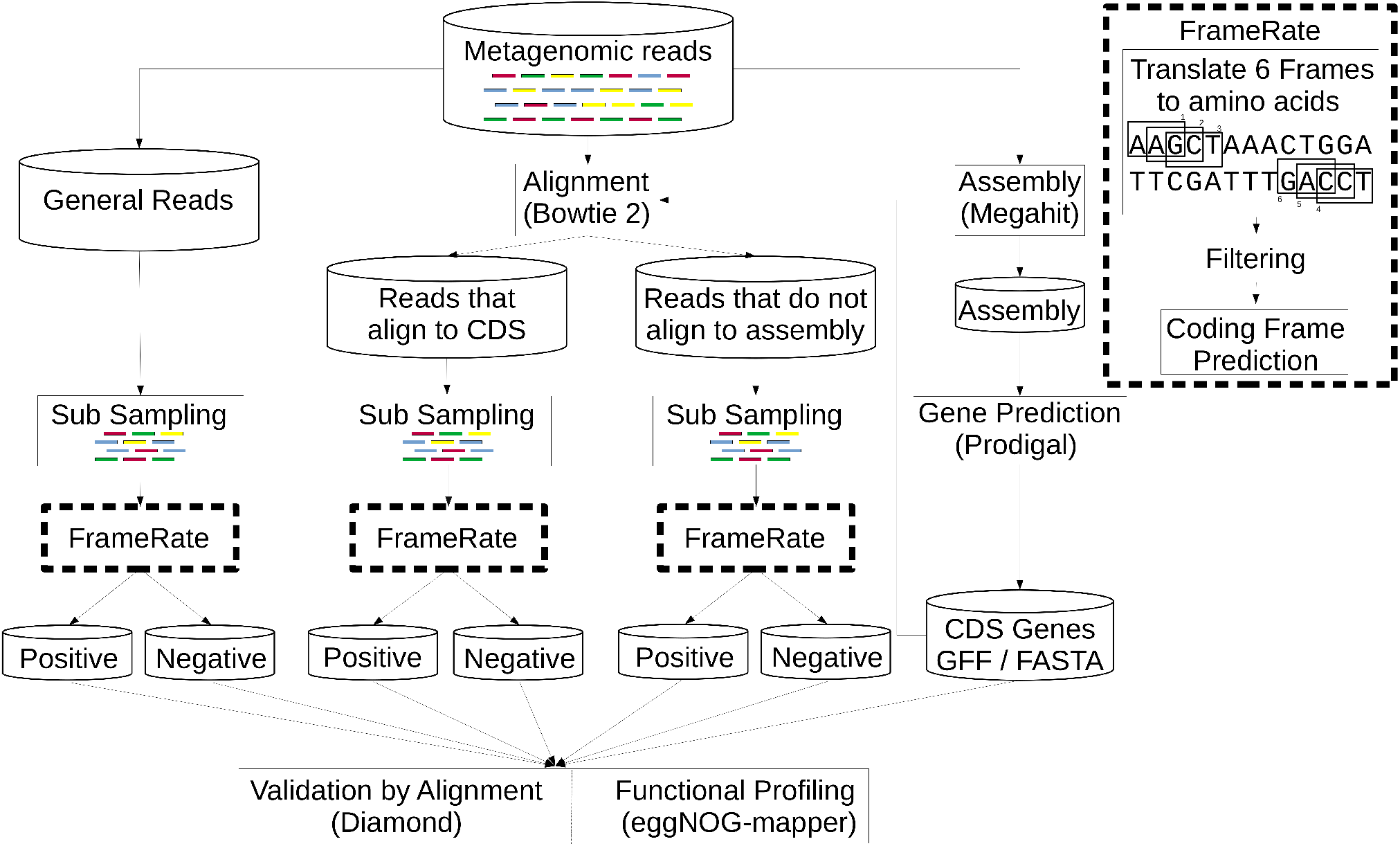
An overview of the applications of the trained FrameRate classifier. The EggNOG COG functional annotations of the Coding and Non-Coding frames classified by FrameRate for the three collections of reads are compared to those identified from the ‘traditional’ CDS gene predictions by Prodigal undertaken on the metagenomic assembly.

### 2.6 Rumen metagenome, Swiss-Prot and Hungate collection protein sequence alignment

The coding and non-coding frames from the the sets of reads (CDS Aligned, Unaligned and General) were aligned to the Prodigal CDS protein sequences from the rumen metagenome assembly, the Hungate genome collection [Seshadri et al., 2018] (a set of 1,436,647 protein sequences from 493 cultured bacteria and archaea genomes that have been independently isolated from ruminant organisms), and the Swiss-Prot curated protein database of 568,002 sequences (downloaded 2022.09.02). The alignment was performed using DIAMOND (-blastp option with default parameters and a minimum bit score of 60).

These three alignments would each report a different validation of the coding and non-coding frames. First, the alignment to the rumen metagenome protein sequences would confirm whether the frames being predicted are both correct (coding or non-coding) and whether they represent true fragments of the complete protein sequences, thus facilitating their use to perform functional profiling. Second, the alignment to the protein sequences from the Hungate collection, predicted by Prodigal, would report the prospect of the classified frames being able to be aligned to a dataset that was produced from a genomics workflow, independent of the metagenomic dataset but still from species known to live in the same environment. Third, the Swiss-Prot alignment is an independent validation not related to rumen environment or the eggNOG database.

### 2.7 EggNOG COG functional annotation

eggNOG-mapper was used to perform the functional annotation of both the CDS genes, the FrameRate coding and non-coding frames identified from the rumen metagenomic dataset. eggNOG-mapper uses the similarities that DIAMOND [Buchfink et al., 2021] identifies for each query sequence to compute assignments of Cluster of Orthologous Groups (COG) functional categories [Tatusov et al., 2000]. These COG functions are then used to produce functional profiles for the three sets of sequences.

## 3 Results

### 3.1 FrameRate predicts correct coding frames in unseen gene families

As a tool to identify correct coding frames of potentially coding sequences, FrameRate was first tested for its predictive ability in unseen gene families. The overall prediction accuracy is consistently above 95% at chunk level (75 amino acids) across all genes on a randomly sampled test set throughout 5 independent runs. To examine the feasibility of applying FrameRate on metagenomic samples in which reads (and entire genes) can be shorter than 225 nucleotides (∼ 75 amino acids), we further computed accuracy scores for genes that are between 50-74 amino acids in the test set, as shown in Figure 3. Although FrameRate performs consistently worse as genes become shorter, prediction accuracy for them still maintains at a level above 85%, despite the fact that genes shorter than 75 amino acids only represent less than 5% of the training data.

**Figure 3:**
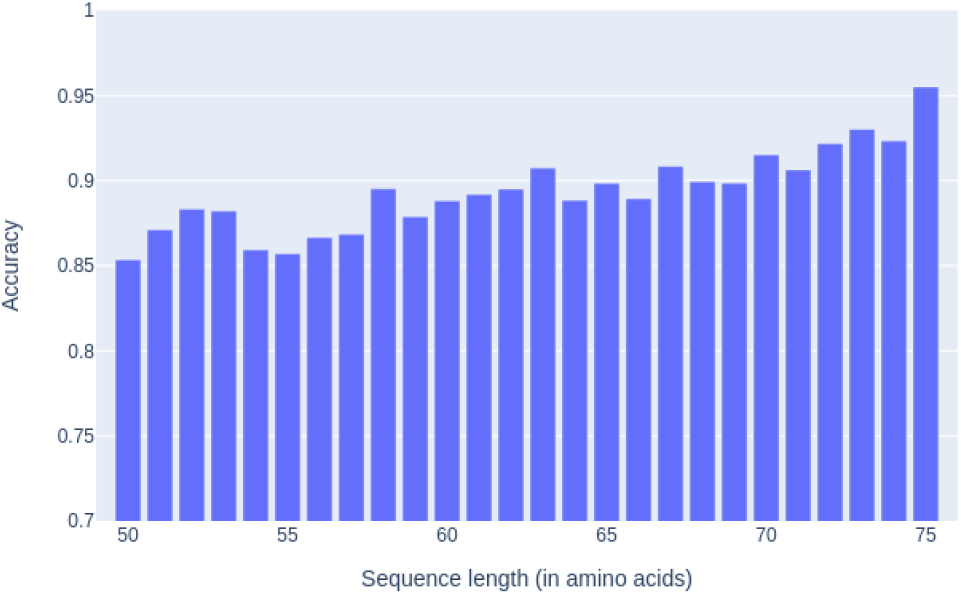
Prediction accuracy by length computed on a balanced test set. The sequences between 50-74 amino acids correspond to full-length short genes which were 0-padded. The bar at 75 amino acids represents the overall accuracy across the entire test set, regardless of length.

### 3.2 FrameRate identifies on average one coding frame per read

Fundamentally, FrameRate was built to identify the correct coding frame(s) from the six possible frames of each read. Since a read could potentially contain more than one coding frame, possibly from overlapping or closely positioned genes, we investigated the proportion of predicted coding and non-coding frames for each read, as shown in Figure 4.

**Figure 4:**
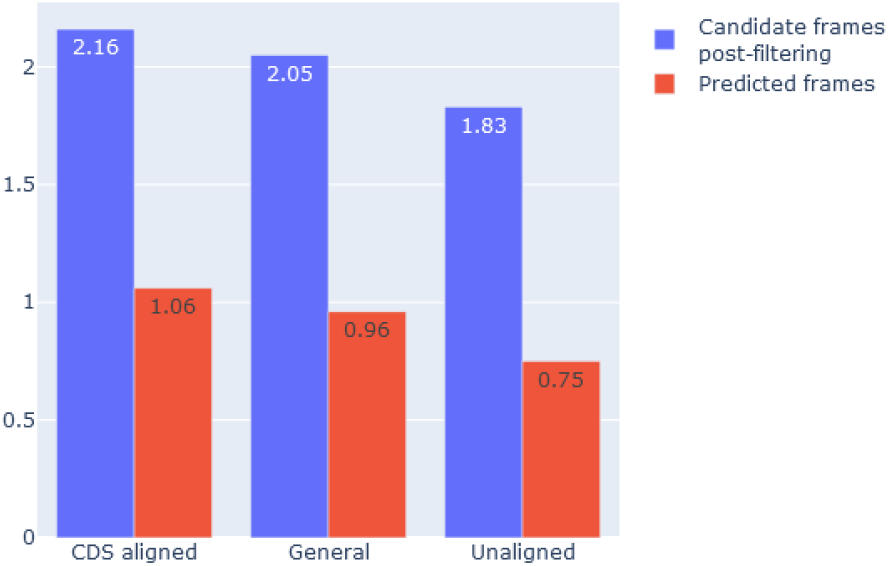
Number of candidate frames (blue) as input to FrameRate vs predicted coding frames (red) as output from FrameRate, computed from the 3 sets of reads.

The majority of the 6 possible frames for the reads in all sets, including those which aligned to the Prodigal CDSs were filtered out, as many negative frames contain internal stop codons (see Subsection 2.3 for further details). Across the 3 sets of reads, only around 2 frames per read passed the stop-codon filtering (and became candidate coding frames as input) and only around 1 frame per read were predicted to be proteincoding by FrameRate. Unaligned reads were reported with the lowest predicted coding frame(s) per read, possibly a result of more of them belonging to the noncoding regions of the genome or being less represented in our training data. Futher details can be found in Supplementary Table 3.

### 3.3 Validation of FrameRate coding predictions through alignment

To investigate whether the predicted coding frames from FrameRate not only represent true fragments of full length protein sequences, but also contain enough information for alignment-based methods (such as those often employed in functional annotation studies), both the coding and non-coding frames were aligned to the following datasets: the proteins translated from the CDS sequences predicted by Prodigal in the same rumen metagenome sample assembly, the Swiss-Prot protein database, and the proteins from the Hungate collection.

As shown in Table 1, there is an extreme disparity between the proportion of FrameRate-predicted coding frames which aligned to each of the three datasets compare to the proportion of predicted non-coding frames. These results suggest that FrameRate is accurately classifying the majority of the coding frames. Nevertheless, the possibility of overlapping genes should be taken into account when analysing the small number of non-coding frames that aligned to either of the 3 datasets. A read spanning two overlapping or closely positioned genes, will potentially contain enough information for FrameRate to classify more than one of its frames as coding at the same time.

**Table 1:** The proportion of FrameRate classified coding and non-coding frames which aligned to the three protein sequence datasets using DIAMOND blastp (protein-protein sequence alignment).

| Dataset | # Protein Sequences | Prodigal Sequences | FrameRate Coding [%] |  | FrameRate Non-Coding [%] |  |
| --- | --- | --- | --- | --- | --- | --- |
|  |  |  | CDS Aligned | Unaligned | CDS Aligned | Unaligned |
| Prodigal CDS genes | 1,647,050 | **N/A** | **80.4%** | **67.3%** | **5.3%** | **8.6%** |
| Hungate Collection | 1,469,083 | **73.0%** | **60.4%** | **51.2%** | **2.8%** | **5.1%** |
| Swiss-Prot | 568,002 | **39.5%** | **23.8%** | **20.7%** | **0.8%** | **1.3%** |

This analysis reaffirms the use of tools such as DIAMOND and eggNOG-mapper (which uses DIAMOND) in this study for aligning sequence data of these short lengths (*∼*75 amino acids) for functional annotation.

### 3.4 FrameRate enables ultra-fast assembly-free functional profiling

Shallow read sampling has previously been used in metagenomic profiling and has been shown to be a successful method of efficiently identifying the majority of the taxonomic profile of a sample [Hillmann et al., 2018]. We compared the functional profiles computed with eggNOG-mapper of the metagenomic Prodigal CDS genes and the 3 read subsamples described above, as presented in Table 2. The functional profiling of the 3 subsamples showed close resemblance to that of the Prodigal CDS genes in two of the EggNOG COG categories, i.e. ‘Cellular Processes & Signaling’ and ‘Information Storage & Processing’. Of note, the functional profiles from the 3 readsets consistently reported a 4% increase for ‘Metabolism’ and a decrease for ‘Poorly Characterized’ compared to the CDS genes, potentially indicating some level of bias or overpresentation either in the Prodigal, eggNOG-mapper or FrameRate methods and annotations. Additionally, there may be differences between the community-derived assembly and individual reads which are being detected by eggNOG-mapper. Therefore, it could be inferred that we are observing not only the above-mentioned functional difference between metagenomically assembled reads (in contig form) and raw read assignment, but also that only a small subsample of an entire metagenomic DNA sample is required to produce a comparative functional profile.

**Table 2:** The EggNOG COG functional categories assigned to the Prodigal CDS genes predicted from the metagenome assembly, the CDS aligned reads, the General reads and the Unaligned reads. Coding Frame (CF), Non-coding Frame (NCF).

| COG Functional Category | Prodigal CDS Genes | FrameRate Coding [%] |  |  |
| --- | --- | --- | --- | --- |
|  |  | CDS Aligned | General | Unaligned |
| CELLULAR PROCESSES & SIG' | **22.26%** | **22.73%** | **22.79%** | **23.15%** |
| INFORMATION STORAGE & PRO' | **21.96%** | **21.21%** | **21.34%** | **21.78%** |
| METABOLISM | **33.23%** | **37.57%** | **37.44%** | **37.02%** |
| POORLY CHARACTERIZED | **22.55%** | **18.49%** | **18.42%** | **18.05%** |

While the majority of coding frames from each read-set were annotated with a COG function, the vast majority of non-coding frames were not (see Supplementary Table 4). Additionally, while the proportion of coding frames with an annotation increased across the Unaligned, General and CDS Aligned datasets, the proportion decreased for the non-coding frames.

While the metagenomic assembly clearly requires access to High Performance Computing (HPC) infrastructure, the FrameRate analysis on a test-set of 10m reads from the same metagenomic sample (the General read set) can be performed on a consumer-grade laptop computer, as shown in Table 3. These reads were computed and classified by FrameRate in 0.5 hours on a system with a GPU and 0.9 hours on a low-powered consumer laptop with 4 CPU cores. The majority of this time was taken up by the IO (input/output) and processing of the reads, such as the DNA to amino acid conversion and stop codon filtering. Moreover, while the entire FrameRate package is less than 10MB in size and only needs to use system memory during runtime, the MEGAHIT assembly also requires substantial disk storage during runtime.

**Table 3:** The time and computation resource requirements of assembly and gene prediction versus FrameRate coding frame prediction. ‘FrameRate 10m’ reports the resources needed to run FrameRate on the set of reads which aligned to the Prodigal CDS genes. The Storage Requirements listed here are the maximum disk space needed during the runtime of each method.

| Analysis | Compute Time<br>[CPU Cores] | Memory<br>Requirements | Storage<br>Requirements |
| --- | --- | --- | --- |
| MEGAHIT Assembly | 46 hours [32] | ~400GB | ~100GB |
| Prodigal CDS Prediction | 0.2 hours [4] | ~2GB | N/A |
| FrameRate 10m Reads<br>GPU | 0.5 hours [1] | ~10GB | 1MB |
| FrameRate 10m Reads<br>CPU | 0.9 hours [4] | ~6GB | 1MB |

## 4 Discussion

### 4.1 Profiling metagenomic samples without assembly

The processes for assembling metagenomic samples involve a number of complex computational methods that each performs a degree of filtering and decisions specific to the tools used. Additionally, the resulting output is often a consensus of the original reads and as such, often does not accurately represent the genomes in the sample [Nicholls et al., 2021]. Therefore, it is important that we continue to develop methodologies that can characterise genomic data in its most raw form. Furthermore, unlike metagenome assembly, the preprocessing conducted with FrameRate is conducted equally on all reads and all potential protein sequences can be classified.

The validation studies of FrameRate produced a clear indication of the ability for the model to differentiate coding from non-coding amino acid sequences. While noting that both the metagenome and Hungate Collection CDSs contain their own limitations, both of these alignment studies not only showcase the high level of FrameRate CFs which have correctly been classified, but also report that its NCFs are significantly less likely to report an alignment to known CDS genes. These results could be used to help filter or align DNA reads without any homology to known genes, reducing the reliance on assembled genomes for identifying coding sequences. The convention is often to ignore these reads and not undertake any further analysis of them. While developments such as using additional rounds of assembly using assembled contigs can help to assemble these unassembled reads, there is still no universally implemented solution [Li et al., 2015, Wick et al., 2017, Hitch and Creevey, 2018].

The coding frames predicted by FrameRate showed a *∼*4% shift from the COG category ‘Poorly Characterized’ to ‘Metabolism’ compared to the CDS genes predicted from the metagenome assembly. This shift was observed in each analysis between FrameRate-classified coding frames and the CDS genes. The consensus sequences produced by the metagenomic assembly, combined with the biases imposed by the CDS gene prediction, are both possible reasons behind this difference. Additionally, FrameRate is able to report each frame if there are two genes positioned over a single read, thus allowing for the reporting of the function from both. If instead an alignment tool is used for functional profiling, not only will this be much slower, but blastx will only return results for a single frame. Even though evidence exists that genes positioned close together (also those overlapping) have been frequently observed to contain similar functions, it is not always the case [Mi-helčić et al., 2019].

### 4.2 FrameRate reduces the resources required for metagenomic profiling

Assembly and annotation techniques are not keeping up with the growing number and size of metagenomic studies being conducted [Lapidus and Korobeynikov, 2021]. As a direct consequence of the vast amount of time and resources required to carry out the analysis, many years can pass by between environmental sampling and the publication of the resulting data and analysis [Haroon et al., 2016, Sunagawa et al., 2020]. An important factor that must be considered in contemporary genomic analysis is the computational resources required to perform each specific step. Another question is whether the potential outcome of the analysis justifies the use of those resources. This is even more true with metagenomic analysis, which most often cannot be performed on local machines.

Machine learning promises a number of benefits for genomics, with relatively small time and computational resource requirements being considered as major factors [Jordan and Mitchell, 2015, Sarker, 2021]. As seen in Table 3, the computation of the metagenome assembly was not only impossible on a personal computer, but also required nearly two days on a HPC with 32 CPU cores and around 400GB of RAM. However, FrameRate was able to report a similar functional profile to the metagenomic CDS genes within a greatly reduced the time-frame and the computational resources of an ordinary desktop/laptop (also note - 4 vs 32 CPU cores required for FrameRate vs MEGAHIT respectively). As FrameRate reported the same functional profile when processing the sampled reads as reported in Table 2, it could be proposed that metagenomic assembly and gene annotation could be bypassed entirely.

## 5 Conclusion

Through the use of FrameRate, we are able to quickly characterise almost every sequencing read, whether it assembles or not, bypassing the biases and limitations existing commonly in assembly-based workflows of metagenomic analysis. While long-read sequencing presents the potential to overcome many of these issues, high-throughput short-read sequencing will likely continue to form the backbone of majority of studies in the foreseeable future.

Furthermore, there are types of genes that while have been identified previously, are still routinely missed by state-of-the-art genome annotation methods and as such are most often missing from canonical annotations of even high quality genome assemblies [Dimonaco et al., 2021]. Therefore, it is important that we continue to develop alternative methodologies that can characterise raw genomic data and supplement existing techniques.

## Author Contributions

All authors discussed the conceptualisation of the model and its application. W.L.W and N.J.D developed the model and code.

