## Supplemental Document for "FrameRate: learning the coding potential of unassembled metagenomic reads"

### Supplementary Information for FrameRate: learning the coding potential of unassembled metagenomic reads

### 1 Supplementary Tables

| | | | | | | | |
|---|---|---|---|---|---|---|---|
| Acetobacter | 14 | Clostridioides | 5 | Methanobacterium | 6 | Salmonella | 235 |
| Achromobacter | 8 | Clostridium | 79 | Methanobrevibacter | 6 | Selenomonas | 7 |
| Acidithiobacillus | 5 | Collimonas | 6 | Methanocaldococcus | 6 | Serratia | 33 |
| Acidovorax | 6 | Corynebacterium | 110 | Methanococcus | 7 | Shewanella | 26 |
| Acinetobacter | 83 | Coxiella | 10 | Methanosarcina | 26 | Shigella | 20 |
| Actinobacillus | 8 | Cronobacter | 10 | Methylobacterium | 11 | Sinorhizobium | 10 |
| Actinomyces | 12 | Cupriavidus | 6 | Microbacterium | 13 | Sphingobium | 9 |
| Aeromicrobium | 7 | Dehalococcoides | 13 | Micromonospora | 5 | Sphingomonas | 9 |
| Aeromonas | 26 | Deinococcus | 11 | Moraxella | 8 | Sphingopyxis | 9 |
| Aggregatibacter | 8 | Desulfitobacterium | 5 | Mycobacterium | 733 | Spiroplasma | 17 |
| Agrobacterium | 8 | Desulfotomaculum | 6 | Mycoplasma | 95 | Staphylococcus | 329 |
| Alcanivorax | 5 | Desulfovibrio | 19 | Myroides | 7 | Stenotrophomonas | 14 |
| Altererythrobacter | 6 | Dickeya | 6 | Neisseria | 36 | Streptococcus | 343 |
| Alteromonas | 19 | Edwardsiella | 10 | Nitrosomonas | 6 | Streptomyces | 82 |
| Amycolatopsis | 9 | Ehrlichia | 15 | Nocardia | 5 | Sulfolobus | 21 |
| Anaplasma | 19 | Enterobacter | 82 | Nostoc | 7 | Synechococcus | 25 |
| Arcobacter | 5 | Enterococcus | 83 | Oenococcus | 5 | Synechocystis | 7 |
| Arthrobacter | 12 | Erwinia | 8 | Paenibacillus | 45 | Thermoanaerobacter | 10 |
| Azospirillum | 6 | Erythrobacter | 5 | Pandoraea | 11 | Thermococcus | 18 |
| Bacillus | 283 | Escherichia | 225 | Pantoea | 14 | Thermotoga | 13 |
| Bacteroides | 17 | Eubacterium | 12 | Paraburkholderia | 9 | Thermus | 8 |
| Bartonella | 36 | Flavobacterium | 16 | Pasteurella | 12 | Thioalkalivibrio | 5 |
| Bdellovibrio | 5 | Francisella | 59 | Pectobacterium | 10 | Treponema | 40 |
| Bibersteinia | 5 | Frankia | 5 | Pediococcus | 6 | Ureaplasma | 10 |
| Bifidobacterium | 79 | Fusobacterium | 15 | Planococcus | 9 | Veillonella | 5 |
| Bordetella | 29 | Gardnerella | 6 | Porphyromonas | 7 | Vibrio | 67 |
| Borrelia | 28 | Geobacillus | 18 | Prevotella | 14 | Weissella | 5 |
| Borreliella | 6 | Geobacter | 11 | Prochlorococcus | 16 | Wolbachia | 8 |
| Brachyspira | 9 | Haemophilus | 39 | Propionibacterium | 26 | Xanthomonas | 57 |
| Bradyrhizobium | 14 | Halomonas | 5 | Proteus | 9 | Xenorhabdus | 7 |
| Brucella | 164 | Helicobacter | 108 | Providencia | 5 | Xylella | 7 |
| Buchnera | 20 | Hymenobacter | 7 | Pseudoalteromonas | 8 | Yersinia | 66 |
| Burkholderia | 183 | Janthinobacterium | 5 | Pseudomonas | 197 | Archaeoglobus | 5 |
| Caldicellulosiruptor | 8 | Klebsiella | 149 | Psychrobacter | 9 | Blattabacterium | 8 |
| Campylobacter | 74 | Lactobacillus | 108 | Pyrobaculum | 8 | Hydrogenobaculum | 5 |
| Candidate | 5 | Lactococcus | 18 | Pyrococcus | 8 | Taylorella | 5 |
| Candidatus | 176 | Legionella | 17 | Ralstonia | 17 | Zymomonas | 7 |
| Caulobacter | 5 | Leifsonia | 8 | Rhizobium | 23 | Myxococcus | 5 |
| Cedecea | 6 | Leptolyngbya | 5 | Rhodobacter | 6 | Hyphomicrobium | 5 |
| Cellulophaga | 5 | Leptospira | 16 | Rhodococcus | 20 | Aerococcus | 6 |
| Chlamydia | 147 | Leuconostoc | 12 | Rhodopseudomonas | 7 | Cyanothece | 6 |
| Chlamydophila | 7 | Listeria | 81 | Rickettsia | 57 | Lysobacter | 5 |
| Chlorobium | 7 | Mannheimia | 15 | Riemerella | 8 | Pseudonocardia | 5 |
| Chryseobacterium | 6 | Marinobacter | 11 | Ruminococcus | 9 | Clavibacter | 5 |
| Citrobacter | 16 | Mesorhizobium | 10 | Saccharomonospora | 6 | | |

Table 1: The 179 Ensembl Bacteria genera with the number of genomes after filtering that were used as training data.

| Data | Number of Seqs | Median Length **[SD]** | Min Length | Max Length |
|---|---|---|---|---|
| Paired Reads | 186,941,580 | 224 **[25.04]** | 38 | 298 |
| MEGAHIT Assembly (Min 1,000bp) | 539,021 | 1,566 **[4,230.01]** | 1,000 | 285,919 |
| Unaligned Reads | 50,412,462 | 218 **[38.02]** | 39 | 298 |
| Metagenome Prodigal CDSs | 1,647,050 | 606 **[630.41]** | 60 | 45,444 |
| Prodigal CDS Aligned Reads | 132,254,283 | 226 **[13.80]** | 39 | 298 |

Table 2: An overview of the sequences (paired reads, contigs or CDSs) for metagenomic data used in this study. (1) This first row describes the raw reads without the input of any metagenome assembly. (2) This row reports the complete set of contigs formed during the metagenomic assembly with a minimum length of 1,000 bp. (3) This row reports the set of raw reads which were not aligned to the metagenomic assembly. (4) This row reports the set of CDS genes predicted by Prodigal from the metagenome contigs. (5) This row reports the reads that aligned to the Prodigal CDS gene sequences which were used later in this study. Standard deviation is abbreviated as **[SD]** and all sequence lengths are reported in nucleotides.

Table 3: The number of coding frames (CFs) predicted by FrameRate for each set of reads, the number of frames remaining for each read after stop-codon filtering, the proportion of CFs classified for each read and lastly the number of non-coding frames.

| Reads (10 million each) | FrameRate CFs | FrameRate Frames Post-Filtering (mean) | FrameRate CFs per Read (mean) | FrameRate NCFs |
|---|---|---|---|---|
| CDS Aligned Reads | 10,563,008 | 2.16 | 1.06 | 11,079,040 |
| General Reads | 9,562,945 | 2.05 | 0.96 | 10,974,171 |
| Unaligned Reads | 7,518,972 | 1.83 | 0.75 | 10,781,585 |

Table 4: Percentage of sequences that have been asssigned to COGs. The sequences predicted to be Coding Frame (CF) have far more COG assignments than the sequences predicted to be Non-Coding Frame (NCF).

| | Percentage seqs assigned to COGs |
|---|---|
| Prodigal CDS Genes | 75.72% |
| CDS Aligned FrameRate CF | 62.58% |
| CDS Aligned FrameRate NCF | 2.99% |
| General FrameRate CF | 60.83% |
| General FrameRate NCF | 4.18% |
| Unaligned FrameRate CF | 55.48% |
| Unaligned FrameRate NCF | 5.89% |

Table 5: Proportion of FrameRate classified coding and non-coding frames which aligned to their respective protein sequence datasets using DIAMOND blastp (protein-protein sequence alignment).

| Dataset | Total CDS Genes | FrameRate Coding [%] | FrameRate Non-Coding [%] |
|---|---|---|---|
| Prodigal CDS genes | 1,647,050 | 8,489,081/10,563,008 **[80.4%]** | 590,511/11,079,040 **[5.3%]** |
| Hungate Collection | 1,469,083 | 6,379,885/10,563,008 **[60.4%]** | 305,351/11,079,040 **[2.8%]** |
| Swiss-Prot Protein Database | 568,002 | 2,513,100/10,563,008 **[23.8%]** | 88,770/11,079,040 **[0.8%]** |

#### 2 Read Trimming

Trimmomatic parameters used to pair-end join the reads from the two trimmed fastq files.

```
java -jar ~/Trimmomatic-0.39/trimmomatic-0.39.jar PE -trimlog \
trim_log.txt SRR873595_1.fastq.gz SRR873595_2.fastq.gz trimmed_paired_SRR873595_1.fastq.gz \
trimmed_unpaired_SRR873595_1.fastq.gz trimmed_paired_SRR873595_2.fastq.gz \
trimmed_unpaired_SRR873595_2.fastq.gz  ILLUMINACLIP:TruSeq3-PE.fa:2:30:10 \
LEADING:3 TRAILING:3 SLIDINGWINDOW:4:15 MINLEN:36 -threads 8
```

#### 3 Gene Clustering with CD-Hit

```
cd-hit -i protein_sequences_to_be_clustered.fa -o clustered_protein_sequences -s 0.8 -c 0.8
-sc 1 -sf 1 -p 1 -g 1 -d 0 -M 10000 -T 8
```

### 4 Classified reads

An example of part of the FASTA output file for the frames that were predicted to be coding by FrameRate. As can be seen even in this small selection, there is a large distribution in confidence scores, ranging from 0.58 to 0.99.

```
>SRR873595:::1167:0:0:0:;0.999013_Frame:4_Score:0.58
ERTHQNAPSLFVPEPKTLHYPPSLPFEEEVEIFFAIQMRKKYHLLLDCLDRLCLAGYTQSGDGRVTCLFLF
>SRR873595:::2486:0:0:0:;0.998817_Frame:1_Score:0.91
YLCTMFVVMMTLFVIVLVGYGAGKLGYLGGDFDRQLSRLVINMTCPALILSSAMTGELPDREYILPLLLISVVTY
>SRR873595:::3064:0:0:0:;0.999312_Frame:5_Score:0.99
INIAGAERYRAITTSHIRNADGAYLVYDITNSSTFENIGFWLETVKKATDDNIVIYLVGNKADLIDSSGRNRRVT
>SRR873595:::4456:0:0:0:;0.986443_Frame:6_Score:0.56
IQSGESDKNGRMVKHGSSALRCVLMRCADSFALHNPVVYEYKLKKMNEGKFFRVALSHVAKKLIRTIYTLEKNDL
>SRR873595:::5428:0:0:0:;0.989339_Frame:2_Score:0.82
LQHPKDIVEGSEAWDAVPDLFLVLVSEASNTSLSPALSLRVYIIYIPVFLSYSLPPFFSFF
>SRR873595:::7972:0:0:0:;0.999263_Frame:2_Score:0.74
IFACRNKTSLMLDRTQTIEKLKSTRQYFSEHYGVSSMLLFGSVARNEQKEDNDVDVCVEMKPNLFKQAGVK
>SRR873595:::9738:0:0:0:;0.999549_Frame:6_Score:0.92
ELEASVSLLDETTSTLEEDRSADSATNVALTSTSKFGISKAQLPSTAVLPIPHIPGFVTLPGSTVVQGTIVLESR
>SRR873595:::9738:0:0:0:;0.999549_Frame:1_Score:0.88
VNRSLDSKTIVPCTTVEPGNVTNPGMWGMGSTAVEGSCAFDMPNFDVEVSATFVAESADLSSSSVEVVSSSSETL
>SRR873595:::9884:0:0:0:;0.987154_Frame:5_Score:0.63
TERRWAV*TALSKTEFRSGEGRGCNLSGSGRYPGNEAVIFDCSELRWHHASSQTNELPSLQQDGSFLFALAPV
>SRR873595:::11158:0:0:0:;0.998169_Frame:6_Score:0.58
KRACA*SWARLSDTPSERRAGWNGLLGSAATQRDPRLPPIGRYGWVQGFLPPALSNIWSNREKRAALVLDRG
```
